# How AlphaFold 3 helps with multiscale drug discovery

**DOI:** 10.64898/2026.10.19.757941

**Authors:** Ratilal Akabari, William Mangione, Zackary Falls, Ram Samudrala

## Abstract

AlphaFold 3 (AF3) predicts protein ligand complexes, but the utility of its confidence scores for drug discovery remains unclear. We benchmarked AF3 interaction scores against BANDOCK+, an experimentally informed approach, using approved drugs and human protein libraries. AF3 ipTM achieved slightly higher observed aggregate disease ranking scores in a matched 1,103 protein comparison, with performance varying among individual indications. BANDOCK+ enabled substantially broader protein coverage using a computationally less demanding approach, including a 20,158 protein reference library. These findings highlight the potential of AF3 confidence scores for computational drug discovery and the tradeoffs between predictive performance and scalability.

---

Accurate compound protein interaction models are essential for large scale drug discovery because interaction quality directly affects compound prioritization and disease ranking. Frameworks such as Computational Analysis of Novel Drug Opportunities (CANDO) use compound protein interaction matrices to perform systematic drug and disease ranking, but experimentally characterized interactions remain incomplete across the human proteome. Computational approaches are therefore needed to expand interaction coverage [1, 2]. Recent advances in artificial intelligence based structure prediction have enabled the generation of protein structures at unprecedented scale. AF3 extends these capabilities by modeling protein ligand complexes and providing confidence measures for predicted interactions [3]. However, accurate structural predictions do not necessarily establish reliable binding affinity estimates or compound ranking performance [4, 5]. Although AF3 confidence metrics have been evaluated for ligand prioritization, their utility for large scale disease ranking and drug repurposing remains less established.

Here, we benchmark AF3 derived interaction scores against Bionalytical docking plus (BANDOCK+), which integrates experimentally characterized interactions from BindingDB with protein and chemical similarity information [6, 7]. The original BANDOCK approach was previously benchmarked against AutoDock Vina within CANDO and achieved higher average indication accuracy using matched protein libraries [8]. Unlike BANDOCK+, AF3 derives interaction confidence from predicted protein ligand complexes. We evaluate both approaches using the same CANDO framework and approved drug and human protein libraries, with ranking performance measured using new average indication accuracy (nAIA) and normalized discounted cumulative gain (NDCG) [9]. We further examine the influence of protein structural quality, model confidence, and disease indication on ranking performance.

We compared AF3 derived interaction scores with BANDOCK+ using a shared experimentally solved human protein library and FDA approved drug library. Among four evaluated AF3 confidence metrics, ipTM achieved the highest overall CANDO ranking performance and was selected for comparison with BANDOCK+ (Supplementary Fig. S1). To assess whether ranking performance varied with protein structural quality or model confidence, we selected protein sublibraries from 5,317 experimentally solved human proteins based on PDB resolution and AF3 base model pLDDT. AF3 and BANDOCK+ showed similar nAIA and NDCG performance in the selected high and low quality sublibraries, whereas BANDOCK+ performed moderately better in the unbiased 200 protein sublibrary (Fig. 1).

**Figure 1:**
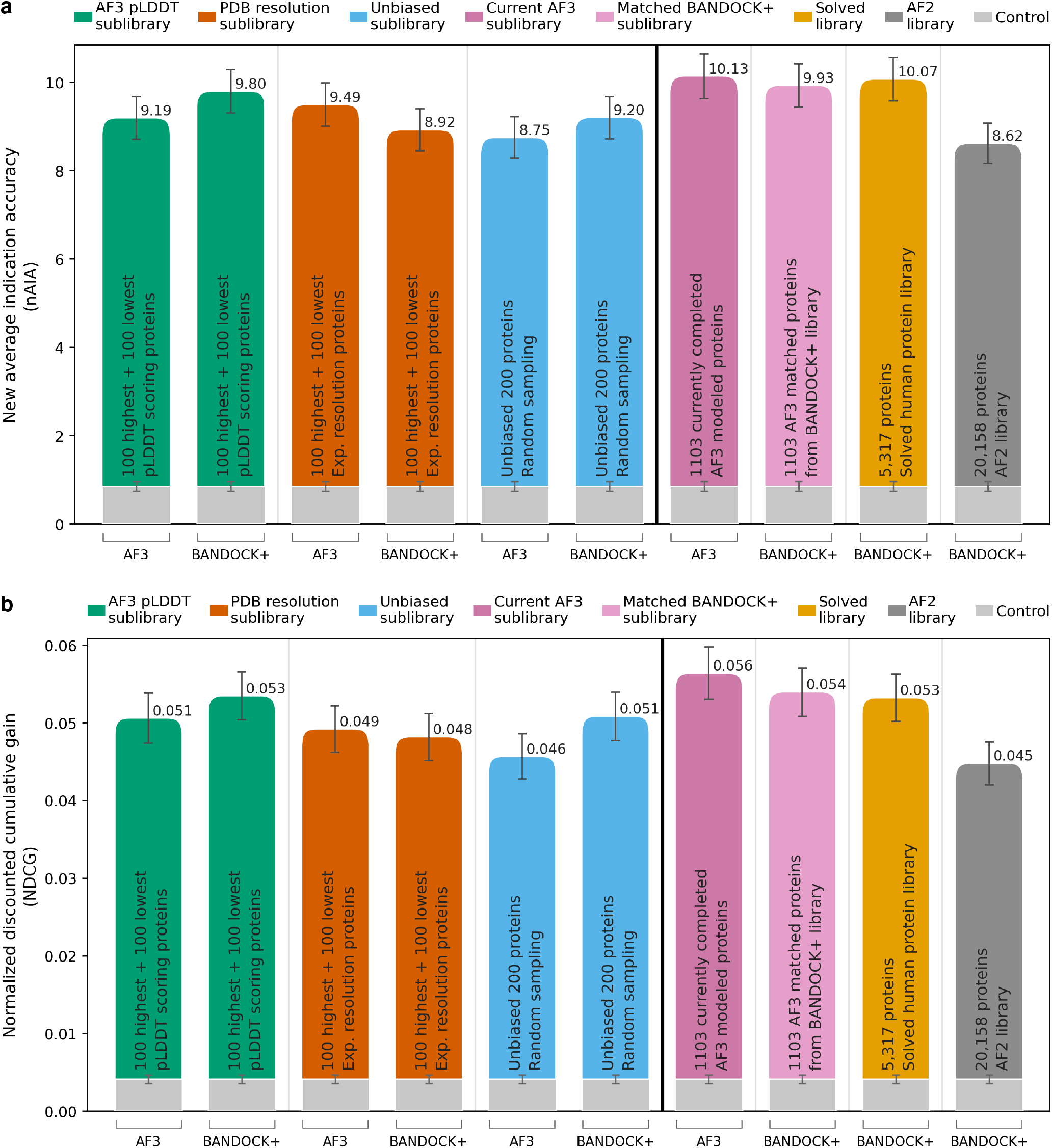
AF3 derived interaction scores achieve comparable compound and disease ranking performance to BANDOCK+. AF3 ipTM scores and BANDOCK+ scores were used to generate compound protein interaction matrices using 2,449 approved drugs and human protein libraries. Protein sublibraries included pLDDT selected, PDB resolution selected, unbiased 200 protein sublibraries, the available 1,103 AF3 modeled proteins, the complete 5,317 protein BANDOCK+ library, and the 20,158 protein AF2 reference library. **a**, nAIA and **b**, NDCG. Error bars represent SEM calculated across individual disease indications. Gray controls represent mean performance from 100 matrices containing uniformly distributed random interaction scores.

As AF3 coverage expanded, we compared interaction matrices from 1,103 AF3 modeled proteins with BANDOCK+ matrices generated for the same proteins and approved drug library. AF3 achieved nAIA of 10.13 and NDCG of 0.056, compared with 9.93 and 0.054 for BANDOCK+, respectively (Fig. 1). AF3 therefore achieved slightly higher observed aggregate scores than BANDOCK+ in the matched comparison, although paired indication level uncertainty estimates did not establish a clear overall advantage for either approach. The complete 5,317 protein BANDOCK+ library and 20,158 protein AF2 reference library were included to provide context for protein coverage and were not used as direct matched comparisons. These results demonstrate the ability of BANDOCK+ to support interaction scoring at substantially greater protein coverage, although increased coverage did not necessarily translate into improved disease ranking performance. Because these reference libraries differ in protein composition and size, their results should not be interpreted as direct matched comparisons with AF3. Evaluation using the original CANDO metrics, AIA and AI-NDCG, also favored AF3 in the matched 1,103 protein cohort, although relative performance varied among the smaller protein sublibraries (Supplementary Fig. S2).

To improve overall accuracy, we used a data fusion pipeline that selected the better performing method, AF3 or BANDOCK+, for each indication in the matched 1,103 protein sublibrary. This analysis achieved nAIA of 12.85 and NDCG of 0.0738, approximately 27% and 31% higher than AF3 alone, respectively. Future work will explore hybrid scoring based on machine learning to further improve disease ranking.

Because aggregate nAIA and NDCG values may obscure differences among disease indications, we examined performance within MeSH disease categories using matched pLDDT, PDB resolution, and unbiased protein sublibraries. AF3 and BANDOCK+ showed broadly similar disease category performance profiles, although the highest performing categories varied by metric and protein sublibrary (Fig. 2). Similar overall patterns were observed using AIA and AI-NDCG (Supplementary Fig. S3).

**Figure 2:**
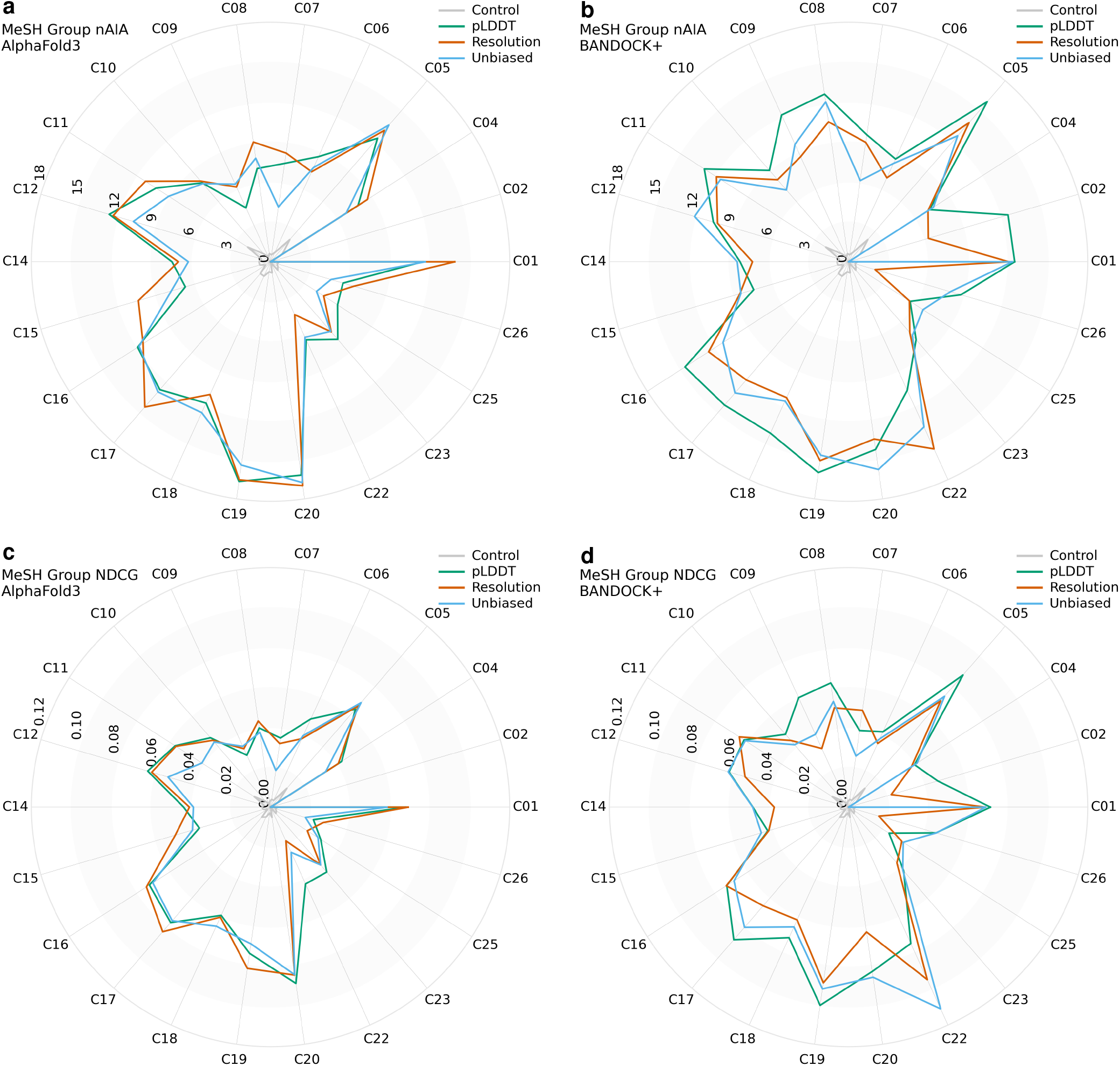
Disease category level comparison of AF3 derived and BANDOCK+ interaction scores. Individual disease indication results from matched pLDDT selected, PDB resolution selected, and unbiased 200 protein sublibraries were grouped by MeSH disease categories. Category means are shown for AF3 nAIA (a), BANDOCK+ nAIA (b), AF3 NDCG (c), and BANDOCK+ NDCG (d). Gray controls represent randomized interaction matrices. MeSH category definitions are provided in Supplementary table. 1

To resolve these indication specific differences, we next compared individual disease indication performance using the matched 1,103 protein libraries. Differences were calculated as BANDOCK+ minus AF3, with positive values favoring BANDOCK+ and negative values favoring AF3 (Fig. 3a,c). Indication level differences favored neither approach consistently. AF3 showed higher indication accuracy for hyperemesis gravidarum, central nervous system tuberculosis, and neurocysticercosis, whereas BANDOCK+ performed better for hyperacusis, subacute bacterial endocarditis, and triple negative breast cancer. Differences in NDCG included higher AF3 performance for alopecia areata and Klebsiella infections, and higher BANDOCK+ performance for migraine without aura and triple negative breast cancer.

**Figure 3:**
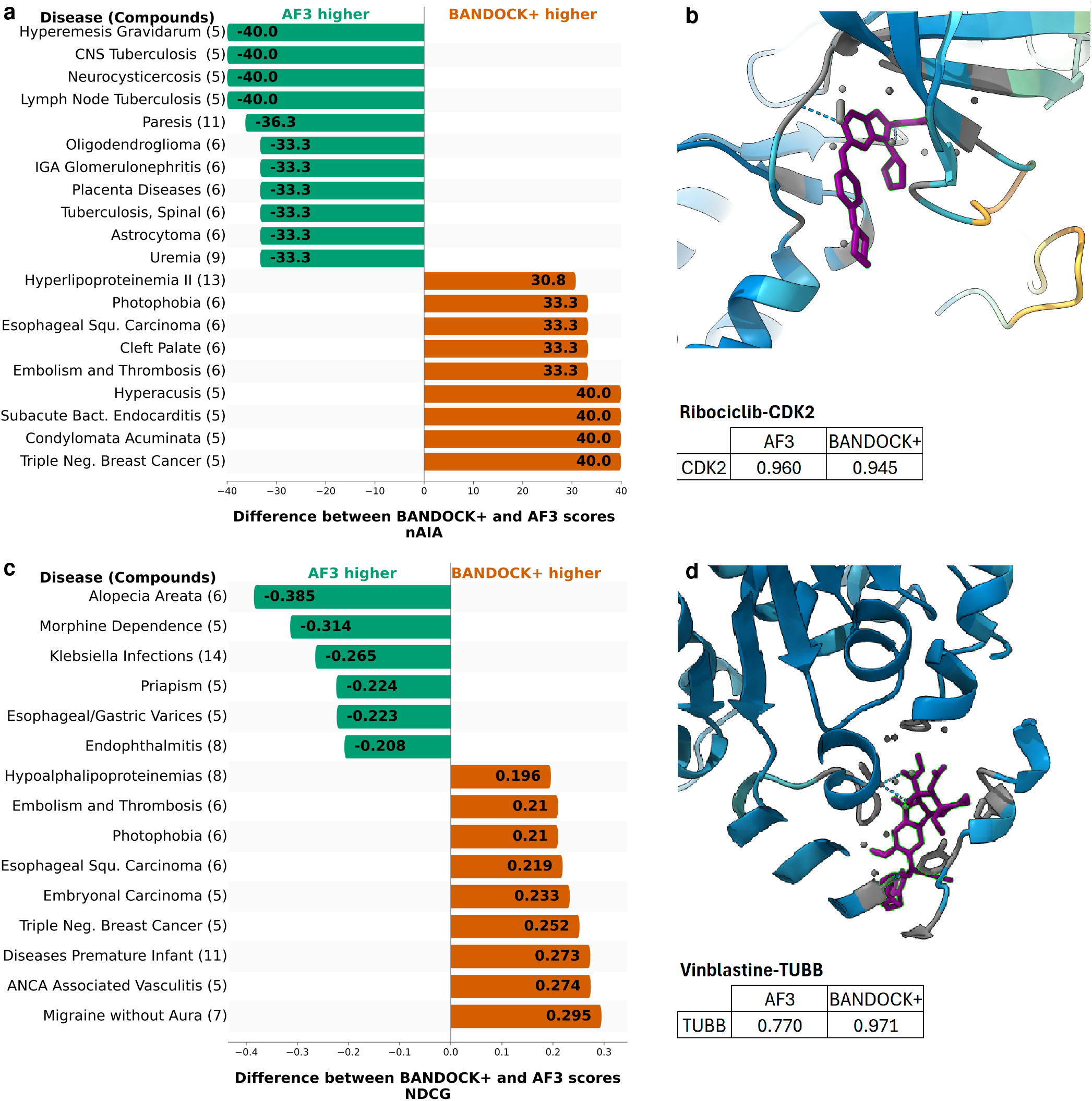
Disease indication level differences between AF3 and BANDOCK+ ranking performance. Differences in indication accuracy **(a)** and normalized discounted cumulative gain **(c)** were calculated as BANDOCK+ minus AF3 using matched 1,103 protein libraries. Negative values favor AF3; positive values favor BANDOCK+. Only indications with at least five associated compounds and extreme differences are included. Representative interactions are shown for ribociclib-CDK2 (**b**; PDB: 4EK4A) in triple negative breast cancer and vinblastine-TUBB (**d**; PDB: 5N5NB) in esophageal squamous carcinoma. AF3 ipTM and BANDOCK+ interaction scores are displayed for each example. These structures represent computational predictions, not experimentally determined protein ligand complexes.

To illustrate the underlying interaction predictions, we examined AF3 predicted complexes for ribociclib-CDK2 and vinblastine-TUBB (Fig. 3b,d). Both approaches assigned high scores to ribociclib-CDK2, while BANDOCK+ assigned a high score to vinblastine-TUBB, consistent with the established tubulin targeting mechanism of vinblastine. These examples illustrate the structural predictions and scoring behavior underlying the disease ranking comparisons.

AF3 interaction confidence scores achieved slightly higher aggregate disease ranking performance than BANDOCK+ in the matched 1,103 protein comparison, although the magnitude of this advantage varied across evaluation metrics and disease indications. This finding is notable given the fundamentally different computational approaches: AF3 generates protein ligand complex predictions using a computationally intensive deep learning model, whereas BANDOCK+ estimates interactions using experimentally informed protein and chemical similarity measures. BANDOCK+ also offers a substantial advantage in computational efficiency and scalability. Importantly, AF3 confidence scores are not direct measures of binding affinity, and the observed disease ranking performance does not establish experimental interaction accuracy or clinical efficacy. These findings demonstrate the potential of AF3 confidence scores for computational drug discovery and highlights the practical advantages of experimentally informed similarity based approaches for large scale screening.

## Methods

### Protein and compound libraries

A library of 5,317 human protein chains with experimentally determined structures available from the Protein Data Bank (PDB) was obtained from the CANDO protein library [2]. Protein sequences were extracted from the corresponding PDB chains and used as inputs for AF3. A previously established library of 20,158 human proteins represented by AlphaFold 2 (AF2) models [10] was evaluated using BANDOCK+ as a reference benchmark for protein coverage and drug ranking performance. All 2,449 compounds were retained without filtering before AF3 modeling. The same compound library was used for AF3 and BANDOCK+ to ensure consistent downstream comparison within the CANDO framework.

### Generation of AF3 interaction scores

AlphaFold version 3.0.1 was used to predict protein ligand complexes for the evaluated human proteins and approved drugs. Complete protein sequences were extracted from individual PDB chains without residue removal, and compounds were represented using their original DrugBank SMILES. Each protein compound pair was modeled using one random seed and five samples. The prediction with the highest AF3 ranking score was selected, and its confidence metrics were obtained from the corresponding summary confidences.json output. Four confidence metrics were evaluated: interface predicted template modeling score (ipTM), predicted template modeling score (pTM), chain pair ipTM, and ranking score. The global iptm value was extracted directly from the selected model’s confidence output and used to construct the primary compound protein interaction matrix. The four confidence metrics were evaluated separately using CANDO, with ipTM achieving the highest overall ranking performance (Supplementary Fig. S1). All 2,449 approved compounds were retained in the interaction matrices for the completed protein cohort, without compound filtering.

### Generation of BANDOCK+ interaction scores

BANDOCK+ interaction scores were generated using experimentally characterized ligand protein interactions from BindingDB combined with protein and chemical similarity measures. Experimental Ki, Kd, IC50, and EC50 values were converted to negative logarithmic molar concentrations, and interactions with measured values of 10 µM or lower were classified as active. UniProt isoform accessions were excluded. Protein structures for BindingDB targets were obtained from the AlphaFold Protein Structure Database (version 6, F1 models). Query proteins were represented by AlphaFold models corresponding to the protein sequences used in the matched AF3 comparisons. Protein similarity was evaluated using glocal sequence alignment with BLOSUM62 (gap opening penalty -11, extension penalty -1, with end gap penalties applied to the query only) and TM score calculated from the same residue correspondences, normalized by query length. The protein similarity score was the mean of TM score and query normalized sequence conservation, defined by identical or BLOSUM62 positive matches. The conservation term was set to zero when the alignment score was negative. Chemical similarity was calculated using Morgan count fingerprints (radius 2, ECFP4 equivalent) and the Dice coefficient. The BANDOCK+ interaction score was the maximum product of protein similarity and chemical similarity among BindingDB ligands with nonzero protein similarity. Scores were generated for the same compound and protein identities evaluated with AF3.

### Benchmarking strategy and protein sublibrary selection

AF3 and BANDOCK+ interaction matrices were evaluated using matched protein sublibraries derived from 5,317 experimentally solved human protein chains. To assess the influence of structural quality and model confidence, proteins were ranked by experimental PDB resolution and AF3 base model pLDDT, respectively. The 100 proteins at each extreme were selected from the full 5,317 protein library for each criterion. Lower PDB resolution values indicated higher experimental structural resolution, whereas higher pLDDT values indicated greater AF3 model confidence. An unbiased sublibrary of 200 proteins was selected by random sampling from the full library. A single random selection was used because of the computational cost of AF3 predictions. The resolution, pLDDT, and unbiased sublibraries were not mutually exclusive, and some proteins appeared in multiple subsets. Identical protein selections were used for AF3 and BANDOCK+ comparisons. The accumulated AF3 cohort comprised 1,103 proteins, including the three predefined 200 protein sublibraries and additional proteins randomly selected from the full library. Overlapping proteins were counted only once. These 1,103 proteins were compared with corresponding BANDOCK+ scores using the same approved drug library. The complete 5,317 protein BANDOCK+ library and the 20,158 protein AF2 based BANDOCK+ library were evaluated separately as reference benchmarks. Neither was treated as a direct matched comparison with AF3.

### Drug discovery evaluation and disease level analysis

AF3 and BANDOCK+ interaction matrices were evaluated using CANDO framework. Identical approved drug libraries, disease associations, ranking procedures, and evaluation metrics were applied to both approaches. The primary metrics were new average indication accuracy (nAIA) and normalized discounted cumulative gain (NDCG), calculated using the revised CANDO benchmarking protocol [9]. Unlike the original average indication accuracy (AIA), which measures recovery of associated drugs within individual compound similarity lists ([11]), nAIA evaluates recovery of withheld drugs from consensus rankings. Additional analyses using the original AIA and average indication normalized discounted cumulative gain (AI-NDCG) are provided in Supplementary Fig. S2. To establish a random control, 100 interaction matrices were generated using values independently sampled from a uniform distribution between 0 and 1. Each matrix matched the protein and compound dimensions of the corresponding interaction library and was evaluated using the same CANDO benchmarking procedure. Control performance was reported as the mean of the 100 evaluations. For disease category analysis, individual indication performance values were assigned to Medical Subject Headings (MeSH) disease categories and averaged within each category. Category specific results were compared between AF3 and BANDOCK+ using matched protein sublibraries. For indication level comparisons, performance differences were calculated by subtracting AF3 values from BANDOCK+ values. Positive differences therefore indicated higher BANDOCK+ performance, whereas negative differences indicated higher AF3 performance. Indications with fewer than five associated compounds were excluded from the waterfall plots.

### Representative compound protein interactions

Representative interactions were selected from disease indications showing differences between AF3 and BANDOCK+ ranking performance. AF3 predicted protein ligand complexes were visualized for ribociclib with CDK2 (protein PDB: 4EK4, chain A) and vinblastine with TUBB (protein PDB: 5N5N, chain B). Corresponding AF3 ipTM and BANDOCK+ interaction scores were retrieved from their respective interaction matrices. The structural representations were used to illustrate predicted complexes associated with disease ranking results and were not considered independent experimental validation of ligand binding.

### Statistical analysis

Ranking performance metrics and their standard errors of the mean (SEM) were obtained from CANDO evaluation outputs. SEM values were calculated from individual disease indication measurements contributing to each averaged metric. Matched comparisons used identical protein subsets and compound libraries. Randomized control performance was summarized as the mean of 100 independently generated random interaction matrices. Differences between AF3 and BANDOCK+ were interpreted descriptively; no formal equivalence testing was performed.

## Data availability

Human protein structures were obtained from the Protein Data Bank (https://www.rcsb.org), and approved compound information was derived from DrugBank (https://go.drugbank.com). Experimentally characterized protein ligand interactions used by BANDOCK+ were obtained from BindingDB (https://www.bindingdb.org). Protein sublibrary definitions, AF3 and BANDOCK+ interaction matrices, compound and protein identifier mappings, and CANDO benchmarking results generated in this study are available on Dryad (https://doi.org/10.5061/dryad.xsj3tx9z4).

## Code availability

Scripts used to prepare AF3 inputs, extract interaction confidence scores, construct compound protein interaction matrices, and perform CANDO benchmarking and visualization are available on GitHub (https://github.com/ram-compbio/alphafold3-drug-discovery).

## Acknowledgements

We gratefully acknowledge use of the research computing resources of the Empire AI Consortium, Inc, with support from Empire State Development of the State of New York, the Simons Foundation, and the Secunda Family Foundation. Support provided by the Center for Computational Research at the University at Buffalo.

## Author contributions

R.S. conceived the initial study. R.S. and Z.F. supervised the research. R.A. developed the AF3 computational workflow, including the AF3 container image, scripts, and analysis pipeline. R.A. performed the AF3 modeling, conducted the computational analyses, interpreted the results, and wrote the manuscript. W.M. developed and executed the BANDOCK+ workflow. R.A. presented the results to the research team and incorporated feedback from all authors into the subsequent analyses and manuscript revisions. All authors reviewed and approved the final manuscript.

## Supplementary information

**Figure S1:**
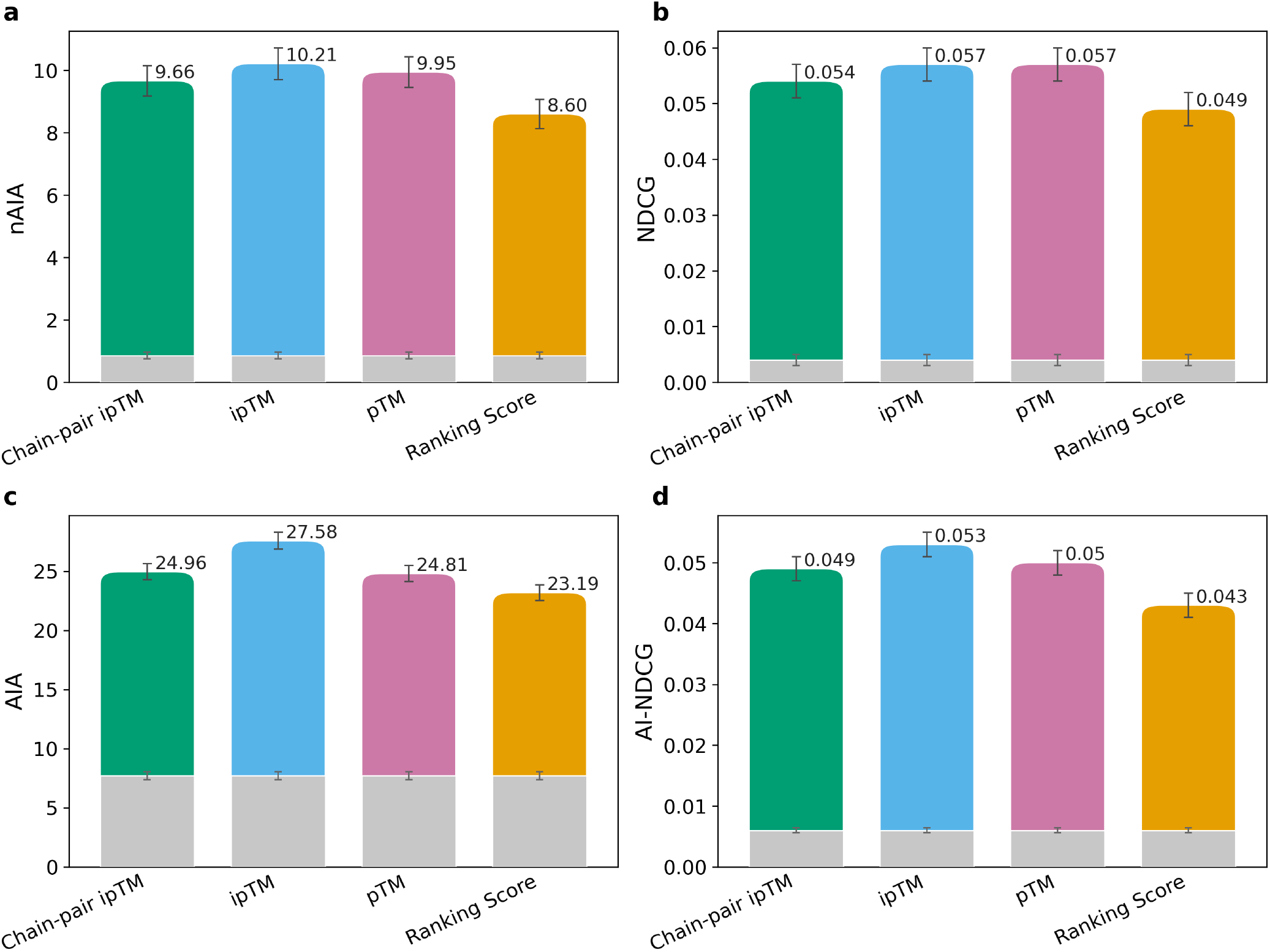
Evaluation of AlphaFold 3 interaction confidence metrics. Comparison of four AF3 derived scores: chain pair ipTM, ipTM, pTM, and ranking score, using CANDO performance metrics: **a**, new average indication accuracy (nAIA). **b**, normalized discounted cumulative gain (NDCG). **c**, average indication accuracy (AIA). **d**, average indication normalized discounted cumulative gain (AI-NDCG) Gray bars represent control values. Error bars indicate the standard error of the mean (SEM). ipTM showed the highest overall performance and was selected for subsequent analyses.

**Figure S2:**
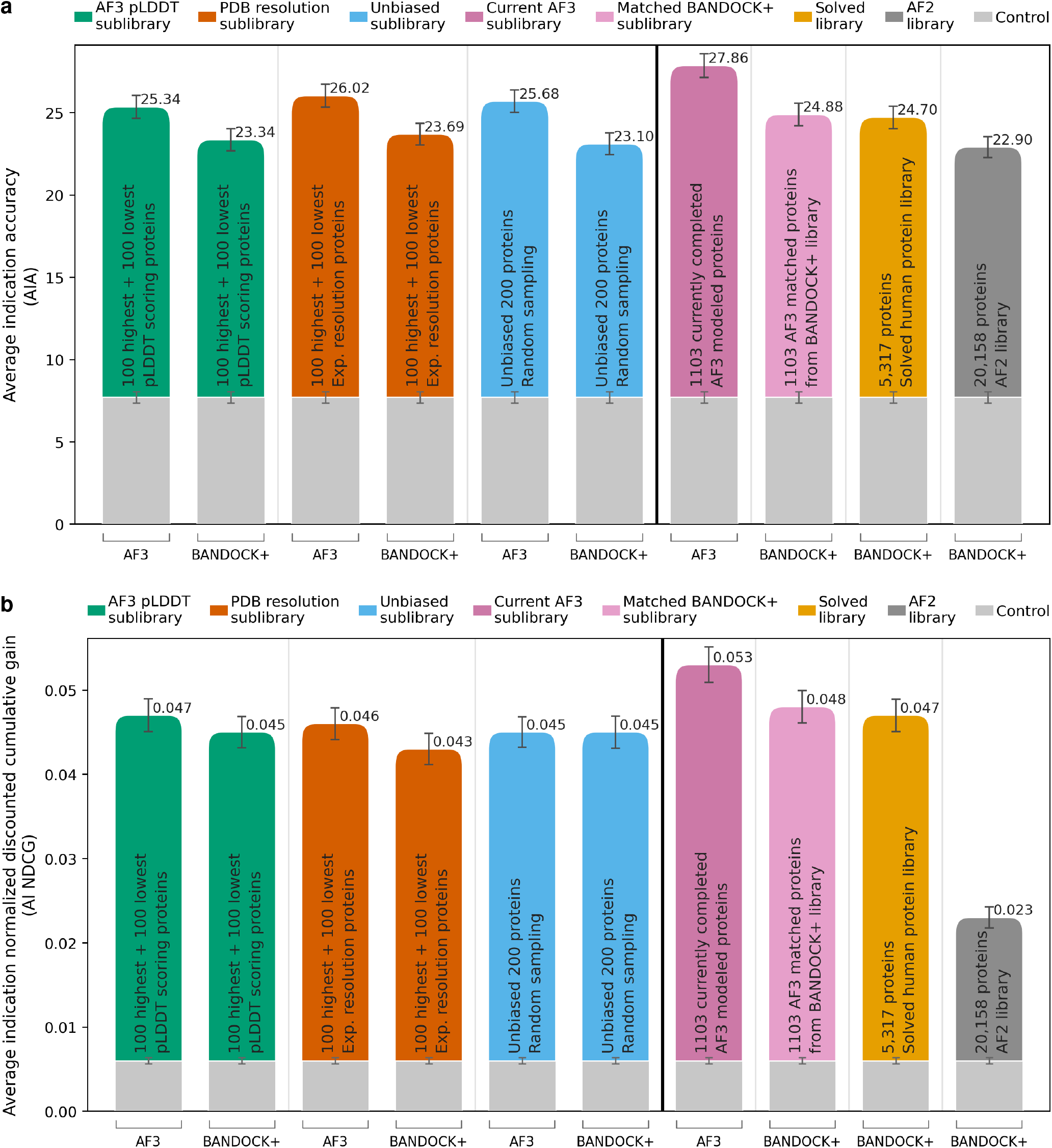
AF3 derived interaction scores achieve comparable compound and disease ranking performance to BANDOCK+ using additional CANDO metrics. AF3 ipTM scores and BANDOCK+ scores were used to generate compound protein interaction matrices using 2,449 approved drugs and human protein libraries. Protein sub-libraries included pLDDT selected, PDB resolution selected, and unbiased 200 protein sublibraries, the available 1,103 AF3 modeled proteins, the complete 5,317 protein BANDOCK+ library, and the 20,158 protein AF2 reference library. a, AIA and b, AI-NDCG. Error bars represent SEM calculated across individual disease indications. Gray controls represent mean performance from 100 matrices containing uniformly distributed random interaction scores.

**Figure S3:**
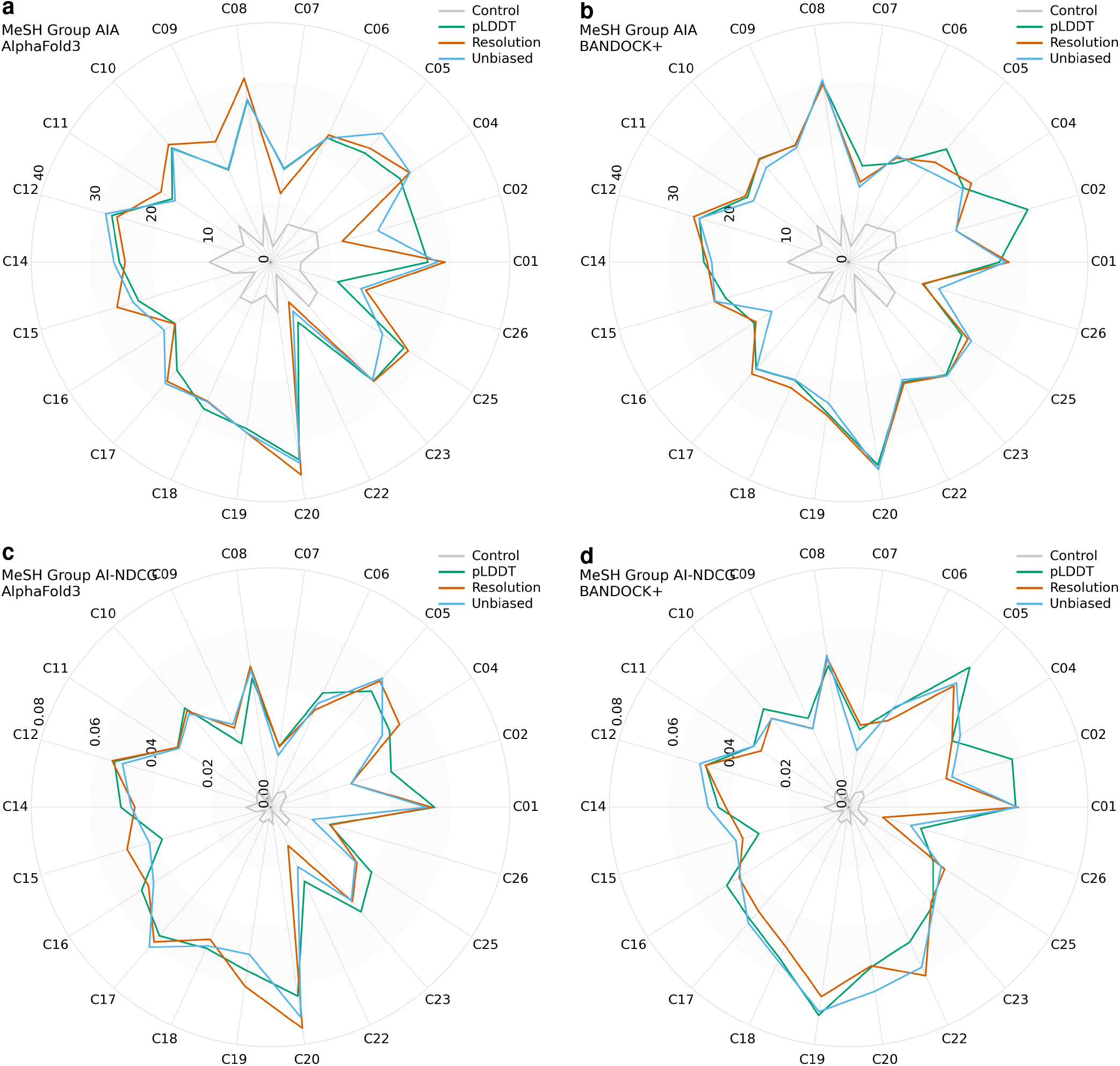
Disease category performance of AF3 and BANDOCK+ using additional CANDO metrics. Individual disease indication results from matched pLDDT selected, PDB resolution selected, and unbiased 200 protein sublibraries were grouped by MeSH disease categories. Category means are shown for AF3 AIA (a), BANDOCK+ AIA (b), AF3 AI-NDCG (c), and BANDOCK+ AI-NDCG (d). Gray controls represent mean performance from 100 matrices containing uniformly distributed random interaction scores. MeSH category definitions are provided in Supplementary table. 1

**Table 1:** MeSH (Medical Subject Headings) categories used for disease ranking evaluation. List of MeSH disease categories and their corresponding classification codes. These categories were used to evaluate and compare disease ranking performance between AF3 and BANDOCK+.

| Code | MeSH disease category |
|---|---|
| C01 | Bacterial Infections and Mycoses |
| C02 | Virus Diseases |
| C04 | Neoplasms |
| C05 | Musculoskeletal Diseases |
| C06 | Digestive System Diseases |
| C07 | Stomatognathic Diseases |
| C08 | Respiratory Tract Diseases |
| C09 | Otorhinolaryngologic Diseases |
| C10 | Nervous System Diseases |
| C11 | Eye Diseases |
| C12 | Urologic and Male Genital Diseases |
| C14 | Cardiovascular Diseases |
| C15 | Hemic and Lymphatic Diseases |
| C16 | Congenital Hereditary and Neonatal Diseases |
| C17 | Skin and Connective Tissue Diseases |
| C18 | Nutritional and Metabolic Diseases |
| C19 | Endocrine System Diseases |
| C20 | Immune System Diseases |
| C22 | Animal Diseases |
| C23 | Pathological Conditions Signs and Symptoms |
| C25 | Chemically Induced Disorders |
| C26 | Wounds and Injuries |

